# Dynamical model of antibiotic responses linking expression of resistance to metabolism explains emergence of heterogeneity during drug exposures

**DOI:** 10.1101/2023.09.22.558994

**Authors:** Mirjana Stevanovic, João Pedro Teuber Carvalho, Philip Bittihn, Daniel Schultz

## Abstract

Antibiotic responses in bacteria are highly dynamic and heterogeneous, with sudden exposure of bacterial colonies to high drug doses resulting in the coexistence of recovered and arrested cells. The dynamics of the response is determined by regulatory circuits controlling the expression of resistance genes, which are in turn modulated by the drug’s action on cell growth and metabolism. Despite advances in understanding gene regulation at the molecular level, we still lack a framework to describe how feedback mechanisms resulting from the interdependence between expression of resistance and cell metabolism can amplify naturally occurring noise and create heterogeneity at the population level. To understand how this interplay affects cell survival upon exposure, we constructed a mathematical model of the dynamics of antibiotic responses that links metabolism and regulation of gene expression, based on the tetracycline resistance *tet* operon in *E. coli*. We use this model to interpret measurements of growth and expression of resistance in microfluidic experiments, both in single cells and in biofilms. We also implemented a stochastic model of the drug response, to show that exposure to high drug levels results in large variations of recovery times and heterogeneity at the population level. We show that stochasticity is important to determine how nutrient quality affects cell survival during exposure to high drug concentrations. A quantitative description of how microbes respond to antibiotics in dynamical environments is crucial to understand population-level behaviors such as biofilms and pathogenesis.

## Introduction

Antibiotic responses in bacteria are remarkably dynamic and heterogeneous. Microbes typically carry regulated mechanisms of antibiotic resistance, which are repressed when the drug is absent to avoid the costs associated with expression of resistance genes [1,2]. Therefore, when challenged by antibiotics, the cell has only a small window of time to activate its defenses before gene expression is halted by the drug action. Since expression of resistance genes is subjected to the strong stochasticity inherent to bacterial physiology, induction of the response is not always successful [3–5]. Sudden exposures to antibiotics have been shown to result in phenotypic heterogeneity, diversity of response outcomes in single cells and complex growth patterns at the population level [6–12]. To understand how microbial populations survive antibiotic treatments, we need models of antibiotic responses accounting for the dynamic and heterogeneous nature of antibiotic resistance.

Following antibiotic exposure, expression of resistance genes is controlled not only directly by regulatory mechanisms, but also indirectly by global effects of the drug on cell growth and gene expression, with a growing body of literature linking metabolism to the ability of bacteria to resist antibiotic treatments [13–15]. Accumulation of resistance proteins in the cell interior depends on the resistance gene’s expression rate and on the dilution of cell components due to cell growth, both of which are affected by the cell’s metabolic environment. The presence of antibiotics itself alters the cell’s growth dynamics and allocation of metabolic resources [16], which in turn affect expression and dilution of resistance, and consequently survival upon drug exposure. During drug responses, antibiotics hamper the induction of resistance genes, which leads to further drug accumulation and decreased expression [17]. These metabolism-mediated feedback mechanisms affect the course of antibiotic responses and can potentially amplify stochastic variations resulting in the coexistence of live and arrested cells [18].

Due to their small size, cellular components in bacteria are often present in small numbers and are subject to stochastic fluctuations. In particular, the transcription factors that regulate gene expression are often present on the order of tens of molecules per cell. Therefore, transcriptional regulation greatly increases stochasticity in gene expression, leading to strong variability in expression levels even among genetically identical cells [19,20]. When this variability is high, it can be harnessed as a resistance strategy [21–25]. Bacterial colonies often harbor a subpopulation of cells with high levels of resistance proteins prior to drug exposure, increasing the chances of colony survival (heteroresistance) [26,27]. However, even in the absence of strong pre-existing heterogeneity, subtle variations in the expression of resistance genes can be amplified during antibiotic responses by feedback mechanisms controlling gene expression, diverging the course of responses among single cells, and ultimately leading to phenotypic heterogeneity at the population level. Therefore, isogenic populations growing under homogeneous environmental conditions still show remarkable diversity of outcomes at the single-cell level during antibiotic responses [21,28].

Here, we develop mathematical models of the dynamics of an antibiotic response, incorporating drug effects on cell growth and gene expression, to explain the emergence of heterogeneity during drug exposures. We start with a deterministic model that reproduces the progression of drug accumulation, expression of resistance and cell growth during a drug response. Then we develop a stochastic model to show how noisy dynamics can lead to phenotypic heterogeneity. We base our models on the classical *E. coli* tetracycline resistance *tet* operon, which displays many general characteristics of regulated antibiotic responses [24,29,30]. The *tet* resistance mechanism is tightly regulated, controls a resistance gene that poses a significant cost for the cell, and is not directly regulated by other cellular processes [6,31,32]. Cells carrying the native *tet* operon were shown to coexist in growing and non-growing states upon exposure to a large dose of tetracycline [6]. The *tet* operon consists of two genes, an efflux pump TetA and its repressor TetR. In the absence of tetracycline, TetR represses both TetA expression and its own [33]. TetR has a strong affinity for tetracycline, binding the drug as it enters the cell, which causes a conformational change resulting in loss of affinity for DNA and release of expression of both TetA and TetR. Efflux pump TetA is then rapidly produced, exporting tetracycline out of the cell. As the intracellular tetracycline concentration decreases, TetR resumes repression, avoiding a toxic overexpression of TetA [34]. Since the *tet* mechanism directly senses the intracellular presence of the drug and elicits a fast and strong response, it is an ideal system to study antibiotic response dynamics and heterogeneity.

We describe the response dynamics of a widespread mechanism of regulated antibiotic response, which is broadly applicable to other responses where the cell needs to react quickly to rapidly changing environments [6,13,35], bringing the system out of equilibrium. Since the dynamics described in our model are not particular to antibiotic resistance mechanisms, many insights from this formulation can also be generalized to other transcriptionally repressed cellular responses that both sense and negatively act upon a chemical signal.

## Methods

### Simulation of the deterministic dynamical model

We numerically integrate the system of differential equations using the ode113 function in MATLAB, using values of external drug concentration and nutrient quality as input parameters. The parameter values used in the simulations are summarized in Table S1 and were either estimated from our experimental data [6], or obtained from literature [24,36]. The code for the simulations is available at our GitHub page: https://github.com/schultz-lab/Phys-Biol-2023.

### Simulation of the stochastic model

We simulate this system using the classical Gillespie’s Stochastic Simulation Algorithm (GSSA) [37], which generates a large number of independent trajectories that are used to calculate probability distributions of the system states over time. Instead of using fixed time steps, the Gillespie algorithm calculates two probability distributions: one for the time that is needed for the next reaction event to occur and a second distribution characterizing which of the possible reactions will occur next. By choosing two random numbers from a random number generator, a value from the time distribution and a reaction are chosen. The time is then increased, and the system is updated accordingly. Each realization of this algorithm represents one trajectory. Transient time-dependent probability distributions can be obtained from a sufficiently large number of trajectories with different random seeds.

## Results

### Dynamical Model of the Tetracycline Response

To capture the dynamics of cell responses, we developed a mathematical model based on the main biochemical interactions involved in the *E. coli* tetracycline response (Figure 1a). We integrate drug diffusion and accumulation into the cell, transcriptional regulation, and expression of resistance genes, as well as global effects of the drug on cell growth and gene expression. Tetracycline is a ribosome inhibitor [38], which reduces cell growth [39] and causes the cell to upregulate ribosome production in response [40]. Altered ribosome levels then result in changes in the partition of the proteome [40,41], affecting expression of non-ribosomal genes including antibiotic resistance (described in detail below). Since in our model the cell growth rate is variable, a cell dies when the drug action causes ribosome function to cease, and the growth rate becomes zero. This system is largely governed by two feedback mechanisms: a stabilizing negative feedback provided by transcriptional repression [42– 46], and a positive feedback mediated by the metabolism (growth rate-dependent) that can lead to bistability [18].

**Figure 1.**
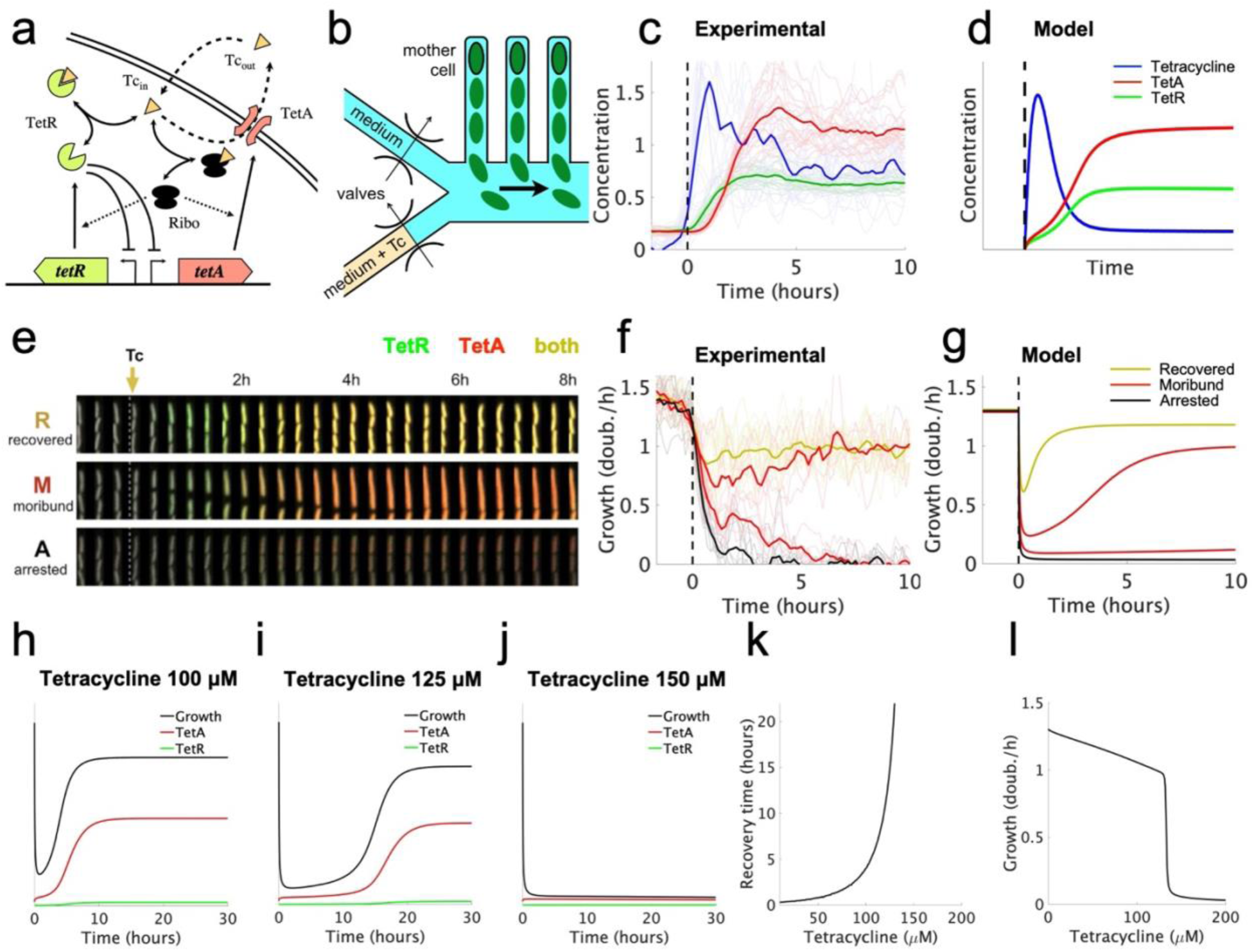
Drug concentration affects recovery from antibiotic exposures. (a) The *tet* resistance mechanism. Tetracycline (Tc) diffuses across the cell membrane, and binds repressor TetR, thereby releasing expression of TetA and TetR. TetA then exports the drug outside the cell. Tetracycline also binds ribosomes, inhibiting protein translation. (b) Schematic of the microfluidic device used for measuring growth and expression of resistance in single cells during drug exposures. The experimental data in this figure is reproduced from [6]. (c) Average tetracycline, TetA, and TetR levels over the course of a response, measured in 40 tetracycline-resistant single *E. coli* cells during a microfluidic experiment. (d) Tetracycline, TetA and TetR levels over the course of a response as predicted by our deterministic model. (e) Three examples of time courses of single cells, resulting in different cell fates. Red and green colors represent expression of TetA and TetR, respectively, measured with fluorescent reporters. (f) Cell growth in single cells following a sudden tetracycline exposure (thin lines). Yellow, red, and black lines correspond to recovered, slow-growing, and arrested cells, respectively. Thick lines represent average growth within each group. Cells that grow slowly following drug exposure eventually either recover or become arrested. (g) Varying the dissociation constant for drug-ribosome binding, the model reproduces the responses seen in single cells. (h-j) Growth rate, TetA, and TetR levels over the course of a tetracycline response, calculated at different extracellular drug concentrations. Higher drug concentrations lead to larger decreases in growth rate, eventually leading to arrest. (k) Recovery times increase with extracellular drug levels, approaching a vertical asymptote at a threshold drug concentration *D*_*thr*_. (l) Growth rate at the end of the tetracycline response for different extracellular drug concentrations. The final growth rate drops sharply around the threshold drug concentration *D*_*thr*_.

The model consists of a system of three differential equations that track changes in repressor TetR (r), efflux pump TetA (*a*), and intracellular drug (*d*) concentrations:

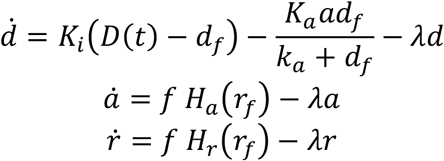

Intracellular drug concentration changes over time according to three processes: the influx of drug from the environment into the cell, the export of drug by the efflux pump TetA, and the dilution of intracellular components due to cell growth. Tetracycline enters the cell by diffusing through the cell membrane, with a rate *K*_*i*_(*D*(*t*) − *df*) proportional to the difference between the extracellular and intracellular drug concentrations (*D*(*t*) and *df*) with a diffusion constant *Ki*. Although diffusion through the hydrophobic membrane is relatively slow, with a half-equilibration time around 45 minutes [47,48], intracellular tetracycline is already toxic for the cell at low concentrations in the nanomolar scale.

Therefore, exposures to extracellular tetracycline in the micromolar scale, which *tet*-equipped *E. coli* can resist, still results in intracellular drug rapidly reaching toxic levels. Efflux pump TetA exports tetracycline out of the cell efficiently, following standard Michaelis-Menten kinetics described by *K*_a_*adf*/(*ka* + *df*), where *K*_*a*_ is the catalytic rate constant, and *k*_*a*_ the Michaelis constant. In growing cells, the intracellular nanomolar concentrations of tetracycline do not significantly saturate TetA. As the cell grows, the drug is diluted in the cell interior, with the dilution rate equal to the growth rate *λd*. The cell growth rate is not fixed, and depends on drug action and metabolism, as detailed below.

In the cytoplasm, tetracycline strongly binds repressor TetR, which then undergoes a conformational change and loses capacity to bind its DNA binding site. Since biochemical binding and unbinding reactions happen at much faster timescales than the other relevant processes, we consider a chemical equilibrium *df* + r_*f*_ *⇌* r_*b*_ between the unbound (free) forms of drug and TetR (*df, r*_*f*_) and the bound form r_*b*_, with an equilibrium constant *K*_*d*_. This equilibrium results in r_*f*_ = r *K*_*d*_/(*df* + *K*_*d*_) and *df* = *d K*_*d*_/(r_*f*_ + *K*_*d*_), and the concentration of free TetR can be calculated by solving the resulting quadratic equation 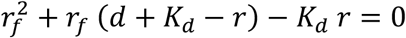. While the bound form of TetR is inactive, the free form transcriptionally regulates expression of both TetA and TetR. TetA concentration changes over time according to its synthesis *f H*_*a*_(*r*_*f*_) and its dilution due to cell growth *λa*. We model TetR regulation of TetA synthesis using a Hill function 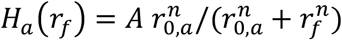, which describes the equilibrium binding of free repressor TetR and its binding sites in the promoter region of TetA. *A* is the fully induced expression rate, in the absence of TetR repression. Free repressor r_*f*_ decreases TetA expression, with *r*_0,*a*_ being the repressor level for half-maximal expression. The Hill coefficient *n* is a measure of how sharply expression rates transition between high and low levels around the threshold *r*_0,*a*_, and is related to the cooperativity in TetR DNA binding. The factor *f*(*λ, d*) modulates the expression of resistance proteins according to global effects of drug action on cell metabolism (cell growth) and is detailed below. TetR concentration is similarly determined by its synthesis *f H*_*r*_(*r*_*f*_) and its dilution due to growth *λr*.

Even in the absence of drug, gene expression depends on the cell growth rate, which is set by the quality of the nutritional composition of the immediate environment. The cell grows according to the total output of translation, and therefore the growth rate is proportional to the ribosomal content of the cell. Faster growing cells then harbor larger ribosomal content, and consequently less non-ribosomal proteins. Additionally, since tetracycline is a ribosome inhibitor, it also causes the cell to upregulate ribosomal content, decreasing non-ribosomal content. A fraction *ϕ*_*Q*_ ≈ 45 % of the proteome, thought to consist of housekeeping genes, rescales gene expression proportionally to the growth rate, balancing synthesis and dilution in order to maintain constant expression levels [40]. To accommodate for changes in ribosomal content, another variable fraction *ϕ*_*P*_ of the proteome adjusts its expression level in the opposite direction. In our model, we account for the effects of intracellular drug concentration and nutrient quality on gene expression and cell growth by incorporating a proteome partitioning framework developed by Scott et al. [40]. According to this framework, all proteins in the cell fall within one of three categories: 1) Sector *Q* proteins whose concentrations are not affected by metabolism (cell growth) or translational inhibition by drug action (e.g., housekeeping genes), 2) Sector *R* of ribosome-affiliated proteins, whose concentration increases with the growth rate or translational inhibition, and 3) Sector *P* of all other proteins, whose concentration decreases with growth rate or translational inhibition to compensate for the increase in ribosomal proteins.

As the fraction *ϕ*_*Q*_ of the cell’s proteome that consists of proteins not affected by translational inhibition does not change, the allocation of resources towards ribosome-affiliated proteins and all others adds to a fixed portion of the proteome, with the two sectors varying according to the cell’s translational (κ_*t*_) and nutritional (κ_*n*_) capacities. κ_*t*_ relates to ribosomal function, measured by the global rate of translation elongation, and κ_*n*_relates to nutrient quality, or the capacity of the culture medium to support growth. Here, we refer to the base value of the cell’s translational capacity in drug-free medium as 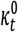, which has a universal value of 4.5 *h*^−1^ for *E. coli* [40] (henceforth, we use subscript or superscript 0 to indicate quantities under full-growth drug-free conditions).

The translational capacity κ_*t*_ is reduced by the presence of intracellular drug, which binds and inactivates ribosomes, and can thus be modeled by 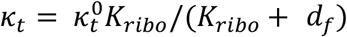, where *K*_r*ibo*_ is the dissociation constant for drug-ribosome binding. Lower *K*_*ribo*_ values correspond to stronger inhibition, and vice versa. The nutritional capacity κ_*n*_ reflects the nutrient quality of the medium, where media with higher κ_*n*_ allow faster growth rates. In situations where nutrient quality does not change, such as in exponential growth in liquid cultures, κ _*n*_ can be determined as a fixed value that fits the maximum growth rate allowed by the culture medium under drug-free conditions (see the SI). Otherwise, decreases in κ_*n*_ can be calculated to reflect nutrient consumption, such as in saturating liquid cultures or as in the nutrient gradients generated by spatial structure in biofilms [7].

According to the framework of the proteome partition, the fraction of the proteome *ϕ*_*R*_ dedicated to ribosomal proteins varies linearly with the cell growth rate, from 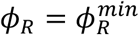 when *λ* = 0 to a theoretical maximum 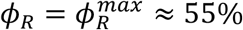, since the remaining *ϕ*_*Q*_ ≈ 45 % of the proteome is fixed and not affected by changes in metabolism [40]. Therefore, the difference between maximal and minimal ribosomal fraction 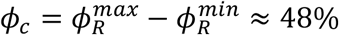 is the variable part of the proteome that can be occupied by the sector *P* proteins that are affected by translation inhibition and nutrient quality. The fractions of the proteome dedicated to ribosomal sector *R* and variable sector *P* proteins are given by 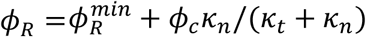, and *ϕ*_*P*_= *ϕ*_*c*_ κ_*t*_/(*κ*_*t*_ + *κ* _*n*_), respectively, such that 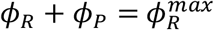. The cell growth rate is proportional to the product of ribosomal content and translation capacity, and is calculated as 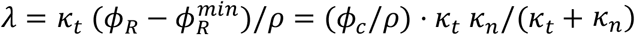, where *ρ* = 0.76 is the conversion factor between RNA/protein ratio and ribosomal fraction calculated for *E. coli*.

Since expression of TetA and TetR has been shown to depend on cell growth, we assume these proteins belong to the variable sector *P*. Therefore, without regulation of the synthesis rates *H*_a_(r_*f*_) and *H*_r_(r_*f*_), we would expect the concentrations of TetA and TetR to scale by *ϕ*_*P*_. We can then calculate the dependence of TetA and TetR synthesis on both translation inhibition and nutrient levels. This dependance is composed of two factors, one reflecting changes in growth rate and another reflecting changes in proteome partition. (*λ*/*λ*_0_) scales down gene expression to match the decrease in growth rate, which does not change expression levels. For proteins in the *P* sector, there is further modulation by a factor 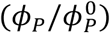, where 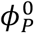 is the *P*-sector fraction under full nutrients and no drug. Therefore, the global metabolic effects on the expression rates of *P*-sector genes can be modeled by 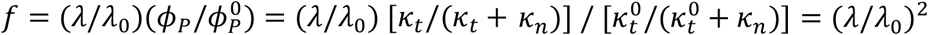. This leads to expression level steady states proportional to ∼κ_*t*_/(κ_*t*_ + κ_*n*_), as expected for proteins in the P sector (see *ϕ*_*P*_ dependence above) and simplifies to *f* = 1 at full nutrients and no drug. We note here that although reducing either the nutritional or translational capacities result in decreased growth rates, they have opposite effects on the proteome partition.

Therefore, while reducing the growth rate by nutrient limitation results in ribosome downregulation and increased TetA and TetR expression levels, reducing the growth rate by tetracycline exposure results in ribosome upregulation and decreased TetA and TetR levels. Next, we use our model to simulate the tetracycline response under different nutrient conditions and drug concentrations.

### Dynamical model captures the dynamics of cell responses

We begin by analyzing the time course of a typical response to a sudden exposure to tetracycline (Figure 1b-g). Initially, as TetA levels are low, the drug quickly diffuses into the cell and accumulates in the cytoplasm, reducing the cell growth rate and slowing down gene expression. As the incoming drug quickly binds and inactivates repressor TetR, expression of both TetR and TetA is released shortly after exposure, although initial accumulation is slow. As TetA levels begin to increase in the cell membrane and export tetracycline back out of the cell, drug accumulation in the cytoplasm slows down and eventually reverses. As intracellular drug levels decline, cell growth and gene expression recover, accelerating TetA accumulation. When intracellular drug returns to low levels, TetR is released, resuming repression. TetA, TetR and tetracycline then equilibrate to steady-state levels, which depend on both TetR regulation and on the effects of the proteome partition on gene expression. This time course of the response dynamics reproduces experimental measurements of expression of resistance genes and cell growth (Figure 1cf), obtained in a resistant population of *E. coli* cells suddenly exposed to a large dose of tetracycline in a microfluidic device [6,49] (data obtained from [6]).

To understand how tetracycline concentration affects cell growth and survival, we simulated the drug response to exposures of increasing drug doses. As drug concentration increases, the growth reduction experienced by the cell at the beginning of a response also increases in both duration and magnitude (Figure 1h-j). At high drug concentrations, synthesis of TetR and TetA is also significantly reduced during this state of translational inhibition, bringing the cell to a quasi-arrested slow growth state. This state can be escaped if the slow production of TetA results in accumulation to sufficient levels to kickstart drug export. With enough TetA, the cell enters a positive feedback loop where reduced intracellular drug concentrations leads to higher growth rates and faster TetA production, resulting in further reduction of intracellular drug levels. As the drug dose increases further, the cell is trapped in the slow growth state for increasingly longer times before eventually recovering. At very high drug doses, however, recovery is no longer possible, with the cell reaching sufficient translational inhibition such that TetA is not produced at high enough rates to initiate a recovery (Figure 1j). The cell then cannot counteract the influx of drug, causing the growth rate to be further reduced by the same positive feedback mechanism.

Increases in drug concentration result in longer recovery times, defined as the time it takes for the cell to recover to the average of its minimum and final growth rates, up to a threshold drug concentration *D*_*thr*_ where the recovery time tends to infinity (Figure 1k). Past this threshold, the cell is permanently arrested following exposure and does not recover. As tetracycline is a bacteriostatic drug, and therefore does not permanently kill the cell (although it can at very high doses [50]) our model considers cell death as a permanently arrested state, that does not lead to recovery unless the drug pressure is lifted. We next determined the stable growth rates at the end of the antibiotic response for different doses of tetracycline, to examine the effect of the drug during steady state. Increasing drug doses cause a reduction in steady-state growth, while still keeping it at relatively high levels up to the *D*_*thr*_ threshold drug dose (Figure 1l). At higher drug doses, steady-state growth is low, corresponding to the arrested state of cells that do not recover. Therefore, the outcome of the tetracycline response is binary, with cells either permanently arrested or recovering to relatively high growth rates, as also observed experimentally. Slow-growing cells are observed only in the transient following drug exposure, and eventually resolve into either arrest or recovery.

Cell survival to drug exposure also depends on the nutrient condition of the culture medium. To determine how growth conditions affect the outcome of antibiotic responses, we simulated the tetracycline response for decreasing values of the nutritional capacity, emulating the nutrient gradients commonly found in structured microbial communities (Figure 2a-d, reproduced from [7]). Decreases in nutritional capacity result in lower growth, both in the presence and in the absence of drug.

**Figure 2.**
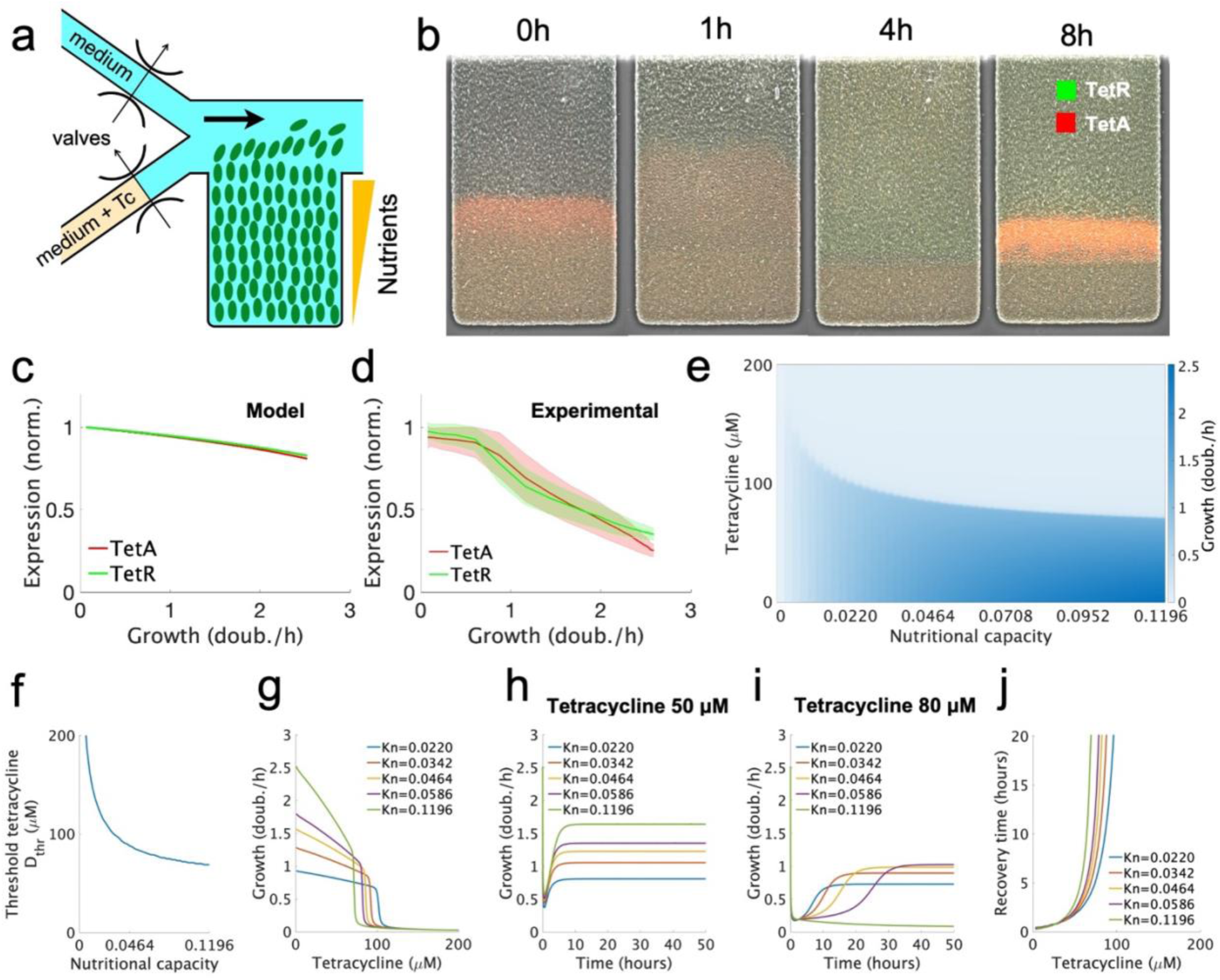
Slow-growing cells survive exposures to higher drug concentrations compared to fast-growing cells. (a) Schematic of the microfluidic device used for measuring growth and resistance expression in a bacterial colony. (b) Expression of resistance genes over time across the colony. The nutrient supply channel (not shown) runs along the top of the trap and flushes away any cells growing out of the trap. The experimental data in this figure is reproduced from [7]. (c) Our tetracycline resistance model, which considers proteome partitioning, predicts a linear decrease in the expression of both resistance proteins with growth rate. (d) Our experimental data also measured a linear reduction in TetA and TetR levels with the growth rate in the absence of tetracycline. (e) Final growth rate at the end of the response for different combinations of extracellular drug concentration and nutritional capacity. Lower nutritional quality allows resistance to higher drug concentrations. (f) Threshold drug concentration *D*_*thr*_ is higher for lower nutritional capacities. (g) Final growth rate varying with extracellular drug concentrations, for different nutritional capacities. (h-i) Growth rate over the course of a tetracycline response for different nutritional capacities, shown at two drug concentrations. (j) Recovery times varying with extracellular drug concentrations, for different nutritional capacities. Cells with lower nutritional capacities have shorter recovery times and maintain growth up to higher drug concentrations.

However, in poor media where the growth rate is low even in the absence of drug, *D*_*thr*_ is significantly higher, which means that cells can recover growth at higher drug doses and show reduced recovery times (Figure 2e-j). Therefore, slow-growing cells in poor media show higher resistance to drug exposures. This result explains cell survival in biofilm microfluidic experiments (Figure 2a-d), where spatial structure causes nutrient gradients decreasing from the surface to the interior of the colony. In such experiments, sudden exposure to tetracycline results in the arrest of fast-growing cells at the surface of the colony, while slow-growing cells in the interior survive exposure and grow to regenerate the surface. Taking the effects of the drug on the proteome partition into account is essential to explain this advantage of slow-growing cells (compare Figure 2e and Figure S1 and the accompanying text in the SI).

Taken together, these results suggests that at high drug doses close to the threshold concentration *D*_*thr*_, small fluctuations could decide the cell’s fate. Since drug responses resolve into either growth recovery or arrest, and both states exist within small margins of drug concentrations and nutrient conditions, we hypothesize that noisy dynamics can lead to the coexistence of live and arrested cells within the same population. Therefore, to study the emergence of heterogeneity during the tetracycline response, we next develop a stochastic model of the response dynamics, which incorporates noise as an integral part of its formulation.

### Stochastic Model of the Tetracycline Response

To simulate the generation of heterogeneity during antibiotic exposures, we developed a stochastic model of the response where the main biochemical reactions are considered as Poisson processes with known rates. The reactions, summarized in Table 1, correspond to the same processes as described by the deterministic model, with drug efflux, transcriptional regulation, and drug action by ribosome binding implemented explicitly. TetA, TetR, intracellular tetracycline, ribosomes and TetR binding sites, both in free and bound forms, are considered as discrete quantities, resulting in a discrete system with known transition rates between states. Stochastic models incorporate noise into their formulation and can be simulated numerically to obtain probability distributions of possible outcomes of the response.

**Table 1.** Reactions of the full stochastic model.

| Reaction | Interpretation |
| --- | --- |
| $\emptyset \xrightleftharpoons{K_i * D} d_f$ | Drug diffusion across the membrane |
| $r_f + d_f \xrightleftharpoons{K_d} r_b$ | Binding of TetR and tetracycline |
| $Ribo_f + d_f \xrightleftharpoons{K_{ribo}} Ribo_b$ | Binding of tetracycline and ribosome |
| $d_f + a_f \xrightleftharpoons{k_a} a_b \xrightarrow{K_a} a_f$ | Tetracycline efflux by TetA |
| $2r_f + O_{1f} \xrightleftharpoons{K_b} O_{1b}$ | TetR binding to $O_1$ site |
| $2r_f + O_{2f} \xrightleftharpoons{K_b} O_{2b}$ | TetR binding to $O_2$ site |
| $O_{1f} f \xrightarrow{f * R/2} O_{1f} + r_f$ | TetR synthesis from $P_{R2}$ |
| $O_{1f} + O_{2f} \xrightarrow{f * R/2} O_{1f} + O_{2f} + r_f$ | TetR synthesis from $P_{R1}$ |
| $O_{1f} + O_{2f} \xrightarrow{f * A} O_{1f} + O_{2f} + a_f$ | TetA synthesis |
| $\emptyset \xrightarrow{g*RR} \mathbf{Ribo}_f$ | Ribosome production |
| $x \xrightarrow{\lambda} \emptyset$<br>$x \in \{\mathbf{Ribo}_f, \mathbf{Ribo}_b, d_f, r_b, r_f, a_f, a_b\}$ | Dilution due to cell growth |

Tetracycline passively diffuses into and out of the cell through the membrane with the same rate *K*_*i*_, resulting in a net diffusion as given by the deterministic model. Inside the cell, tetracycline binds TetR reversibly into an inactive compound. Drug efflux follows Michaelis-Menten dynamics, starting with a reversible binding of tetracycline to TetA, followed by transport of the drug to the extracellular medium. Protein synthesis is combined into a single reaction, with a rate that depends on ribosome availability (detailed below). Repressor TetR regulates protein synthesis by binding to its DNA binding sites and blocking transcription initiation, and therefore TetA and TetR syntheses are modulated by the occupancy of the binding sites. TetR binds two sites *0*_*1*_and *0*_*2*_ as a dimer in the promoter region of the *tet* operon (Figure S2 and the accompanying text in the SI). TetA is synthesized from a single strong promoter *P*_*A*_, which is active only when both binding sites are free. TetR is synthesized from two independent promoters of similar strength: *P*_R1_ is active when both binding sites are free, and *P*_R2_is active when *0*_*2*_ is free. Dilution of cellular components is modeled as a death process, with a rate equal to the cell growth rate *λ*. The reaction rates *K*_*i*_, *K*_*a*_, *R, A*, and *λ* are the same as in the deterministic model (as summarized in Table S1). The forward and backwards rates for the equilibrium constants *K*_*d*_, *K*_*ribo*_, *K*_*b*_ and *k*_*a*_were chosen based on typical values for the timescale of each reaction: binding of small molecules to proteins is in the order of milliseconds in bacteria, while binding of transcription factors to DNA is in the order of seconds.

To incorporate the effects of drug action and nutrient quality, we explicitly consider a ribosome pool of variable size *N*_*R*_, which changes according to the nutritional and translational capacities of the cell. We set a theoretical maximum size of the pool 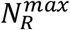 corresponding to the maximum ribosomal content 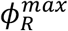. While it is impractical to set 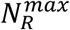 to typical numbers of ribosomes per cell (tens of thousands), a reduced pool still captures the effects of ribosome binding by the drug on protein synthesis, cell growth and proteome partition. Then, the proteome partition is calculated from the size of the ribosome pool, with 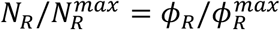. Correspondingly, the P sector fraction can be calculated from 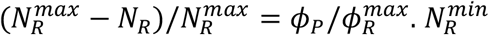 is the minimum ribosome content when growth is zero, corresponding to 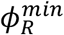, and can be obtained from 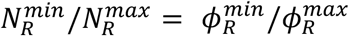, which yields 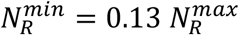. We can therefore calculate the cell growth rate as 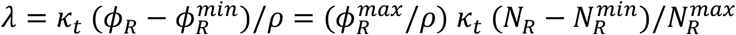. Tetracycline binds and inactivates ribosome, with an equilibrium constant calculated from the drug concentration necessary for half-repression of cell growth, determined experimentally. The reduction in translation capacity then reflects the inactivation of a fraction of the as 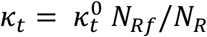 being the number of free ribosomes.

The theory of proteome partition assumes that ribosome production is regulated to achieve optimal levels in each nutrient and drug condition, with the resulting proteome partition affecting the expression of proteins in the P sector. Therefore, we take the approach of calculating the ribosome production rate based on the translational and nutritional capacities κ_*t*_ and κ _*n*_, and then using the resulting changes in the size and availability of the ribosome pool to calculate the effects of drug action in the growth rate and expression of TetA and TetR. In the absence of drug, the average size of the ribosome pool is determined by the nutritional capacity of the medium and the maximal translational capacity as 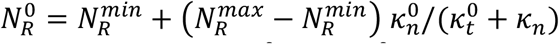. Therefore, we define a basal rate of ribosome production 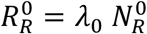 to generate a pool of the expected size. Finally, we calculate the dependencies 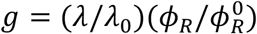 and 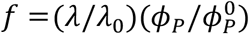 that modulate the synthesis of ribosomes and TetA/TetR under drug exposure, respectively. These dependencies have a common factor 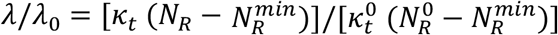, which adjusts gene expression to the reduced growth rate without changing expression levels. Additionally, ribosome synthesis is modulated by 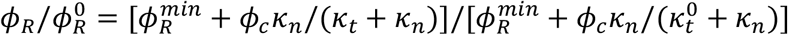, reflecting the upregulation of ribosomes when translation is compromised, and TetA/TetR synthesis is modulated by 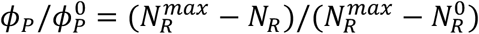 reflecting changes in the proteome partition. We note that the factor *λ*/*λ*_0_ that affects ribosome production is calculated with the growth rate extracted from the pool itself, which is also used to calculate the dilution rate, and therefore does not affect the expected pool size. Since we calculate 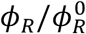 using κ_*t*_ and κ _*n*_ instead of fractions calculated from the pool, the expected pool size does not depend on itself. However, 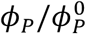 is calculated from the pool, so that gene expression responds to changes in the pool size.

### Stochastic model shows large variations in response dynamics

We started by simulating the system during a drug response to moderate drug concentrations, first in the absence of drug for a period of 1000 minutes, and then introduced an extracellular drug concentration of 100 μM (Figure 3 a-e). The stochastic model closely follows the TetA, TetR and intracellular drug trajectories observed in the deterministic model following drug exposure, with expression rates depending on transient changes in partition fractions and ribosomes. Both in the absence and presence of drug, the size of the ribosome pool equilibrates to the expected steadystate levels. The high influx of tetracycline into the cell following exposure is counteracted by a temporary upregulation of ribosome production, which maintains protein expression, but reduces the resources available to P-sector proteins (TetA and TetR).

**Figure 3.**
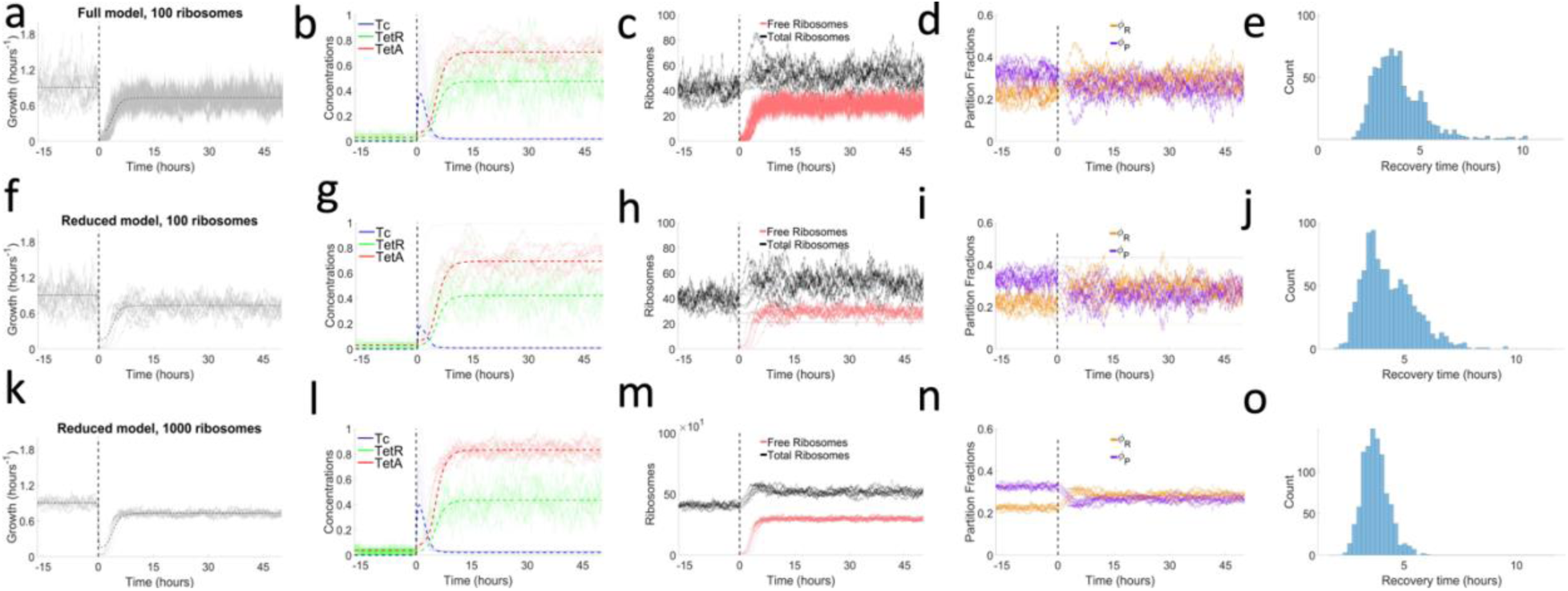
Stochasticity during antibiotic responses to moderate drug concentrations. Time courses of responses to a sudden drug exposure at time 0. Vertical dashed lines indicate drug exposure. (a-e) Full stochastic model, with tetracycline binding reactions considered explicitly, and a pool of 100 ribosomes. (a) Cell growth during the drug response. (b) Intracellular tetracycline, TetR, and TetA concentrations, normalized to maximum values. Thin lines show different trajectories obtained from the stochastic model. Dashed lines indicate the corresponding concentrations obtained from the deterministic model. (c) Ribosome pool, showing total and free ribosomes. (d) P- and R-sector fractions of the proteome. (e) Distribution of recovery times obtained from simulating ∼1000 trajectories. (f-j) Reduced stochastic model, considering tetracycline binding reactions in chemical equilibrium, and a pool of 100 ribosomes. This approximation maintains similar qualitative behavior and dynamics while being simulated at much lower computational cost. (k-o) Reduced stochastic model and a pool of 1000 ribosomes. An increased ribosome pool results in decreased noise and a narrower distribution of recovery times.

The full model, despite being exact, is computationally costly due to the high number of reactions being simulated. To generate a large number of trajectories efficiently, we identify the fastest reactions in the system (Figure S4), which contribute less noise, and approximate their effect using their adiabatic limit. We find that most computational power is spent on fast binding/unbinding reactions of tetracycline with its ligands.

Eliminating explicit tetracycline binding greatly improves computational speed while keeping the most significant sources of stochasticity. The system then shows stochastic noise coming only from the slower synthesis/degradation, drug import/export and DNA binding reactions.

Free tetracycline molecules reversibly bind TetR, TetA and ribosome, with equilibrium constants *K*_*d*_, *k*_*a*_and *K*_r*ibo*_, respectively. Therefore, the copy number of the free and bound forms can be estimated from the total number, such that *r*_*f*_ = *r*_*T*_*K*_*d*_/(*d*_*f*_ + *K*_*d*_) is the number of free TetR molecules, *a*_*b*_= *a*_*T*_*d*_*f*_/(*d*_*f*_ + *k*_*a*_) is the number of bound TetA molecules and *Ribo*_*f*_= *N*_*r*_ *K*_*rib&o*_/(*K*_*ribo*_+ *d*_*f*_) is the number of free ribosomes. Since the export of tetracycline out of the cell is proportional to the number of bound TetA complexes, we arrive at an export rate of *K*_*a*_a_*T*_*d*_*f*_/(*k*_*a*_+ *d*_*f*_) as in the deterministic model. Finally, we calculate the number of free tetracycline molecules from 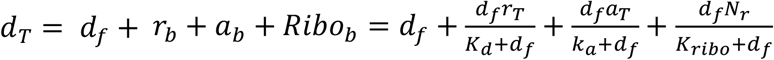, where *b* denotes the bound forms. This equation is solved numerically to obtain *d*_*f*_ at each time step in the simulation.

In this reduced stochastic model, only total copy numbers of the chemical species are modeled explicitly, while free and bound forms are calculated from the equilibrium constants and used to calculate the remaining reaction rates. Simulations performed for the reduced model showed that the overall response dynamics closely followed the full model, and still displayed strong variability of recovery times. We tested ribosome pool sizes of 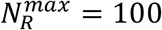 (Figure 3f-j) and 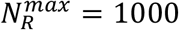 (Figure 3k-o). Although the reduction of the ribosome pool introduces noise into the system, drug responses show qualitatively similar behavior in either case. The size of the ribosome pool is therefore a key parameter measuring the amount of stochasticity introduced into the system by the drug action on cell growth and gene expression. These results suggest that such adiabatic approximations are valid, and the reduced model can be used to generate probability distributions of response outcomes.

### Exposure to high drug doses results in coexistence of growing and arrested cells

We now use the reduced model to simulate a large number of trajectories of the system (∼1000) to obtain a probabilistic distribution of the system’s behavior. In each simulation, the system starts with the expected concentrations in the absence of drug, calculated from the deterministic model, and is simulated for 1000 minutes to reach equilibrium before drug is introduced. The system is then simulated in the presence of drug until steady-state dynamics are reached. In multistable systems, stochasticity can make the response reach different stable points. Therefore, trajectories starting from the same initial conditions can take increasingly different paths, resulting in either recovery or arrest and showing widely different recovery times. Within an isogenic microbial population in a homogeneous environment, different trajectories can be seen as the fates of individual cells, with the probability distribution of response outcomes reflecting the generation of heterogeneity in the population during drug responses. For each drug and nutrient condition used in the simulations, we calculate the probability that responses will result in growth recovery, and, from the subset of recovery trajectories, we calculate the distribution of recovery times.

Simulations of the stochastic model during exposure to high drug doses result in large variations in intracellular drug accumulation, eventually leading to the coexistence of recovered and arrested trajectories (cells). This regime, which includes short recoveries and cell arrest (infinitely long recoveries), corresponds to the very long recovery times observed in deterministic simulations when the drug concentration approaches *D*_*thr*_(Figure 4a). While in some trajectories the cell was able to curb the influx of drug relatively quickly, in others intracellular drug reached much higher levels. Recovery times strongly depend on the maximum drug levels reached inside the cell. When intracellular drug reaches high levels, expression of TetA is compromised, and the cell becomes trapped in a slow-growth state. Taken together, these results suggest the existence of a semi-stable low-growth state, from where the system can escape to recovery if TetA concentrations are high enough. When intracellular drug levels are kept to lower levels, recovery is much quicker. Therefore, the initial dynamics of the response is subject to strong fluctuations, and it is crucial to determine cell fate.

**Figure 4.**
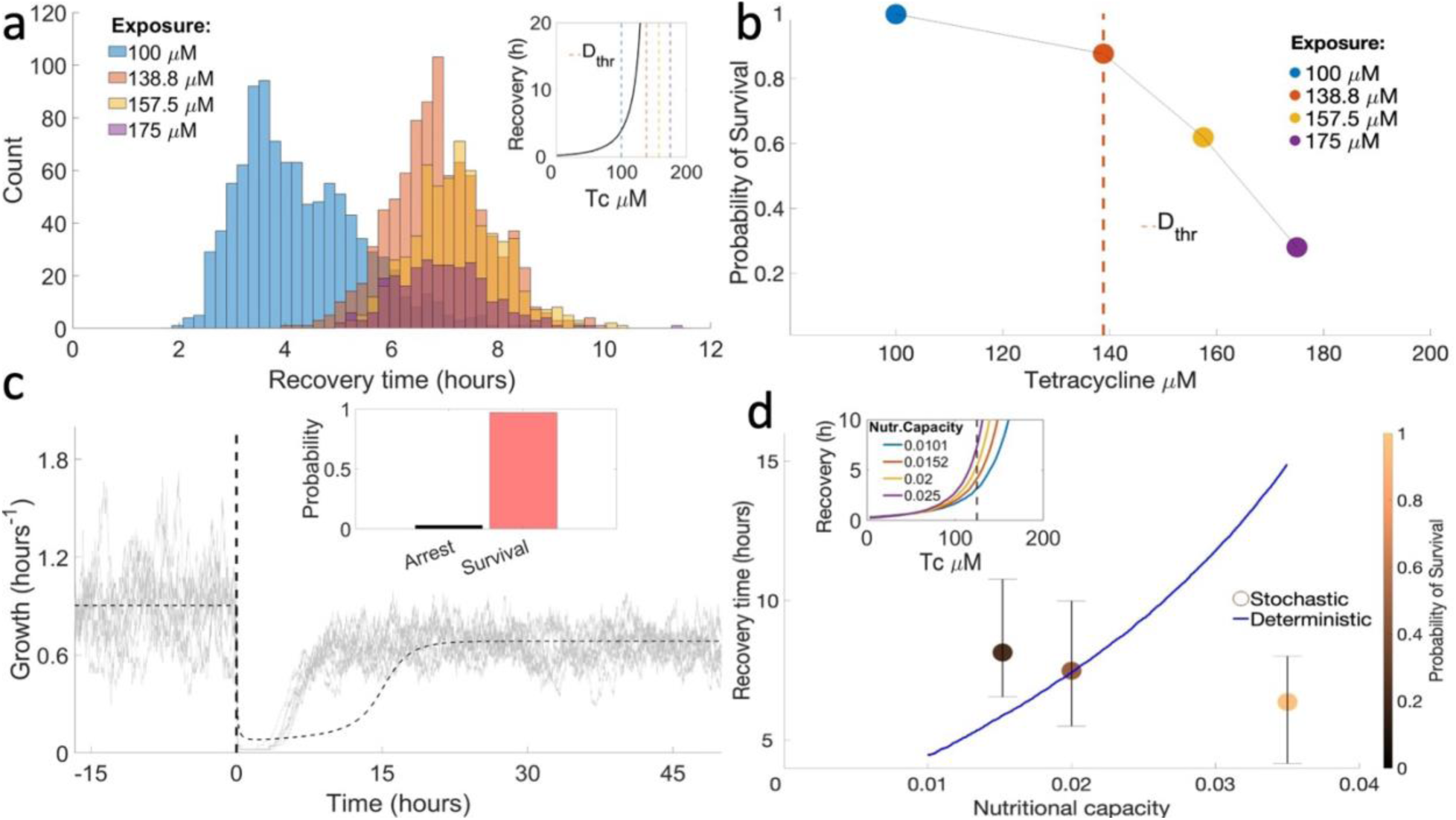
Drug concentration and nutrient quality affect survival rates and recovery times upon exposure to high concentrations of tetracycline. (a) Distribution of recovery times among recovered trajectories, obtained from 1000 simulations for each drug concentration. The mean recovery time increases with extracellular drug concentration up to ∼ 7 hours, obtained when drug concentration reaches the threshold *D*_*thr*_. Increasing drug further does not change recovery times, but results in decreased survival. Inset: Recovery times obtained from the deterministic model, with dashed lines representing the drug concentrations used in the stochastic simulations. (b) Probability of cell survival for different drug concentrations. Survival probability decreases with the drug dose and falls sharply for drug concentrations past *D*_*thr*_. (c) Growth rate in different trajectories during an exposure to 125 μM tetracycline (dashed vertical line). Stochastic fluctuations allow TetA to reach threshold levels necessary for recovery faster than in the deterministic model, resulting in faster recovery. Inset: Probability of survival for this set of simulations. (d) Recovery times for varying nutritional capacities, in the deterministic and stochastic models. The color gradient shows the probability of survival in simulations of the stochastic model. Inset: Recovery times obtained from the deterministic model for different nutritional capacities. The dashed line represents the 125 μM tetracycline concentration used in the stochastic simulations.

At very high drug doses, in regimes where deterministic simulations no longer recover, stochastic simulations may still result in growth recovery, although they become increasingly rare and recovery times tend towards higher values. For exposures to drug concentrations past *D*_*thr*_, we no longer see an increase in recovery times, with distributions still centered at ∼7 hours. However, larger drug levels result in decreased survival, and at very high drug concentrations no cells survive (Figure 4b). These results agree with the experimental observation that cell fate is decided within the first few hours after exposure, after which recovery is rare.

Next, we investigated how nutrient conditions affect the dynamics of the antibiotic response during an exposure to a moderate drug concentration. For high nutritional capacities, average recovery times are faster than those predicted by the deterministic model (Figure 4c). Since recovery depends on TetA levels reaching high enough levels to reverse the influx of drug, stochasticity in TetA expression increases the probability that this threshold is reached earlier. However, unlike in the deterministic model, lowering the nutritional capacity does not result in shorter recoveries (Figure 4d).

Although the probability of recovery decreases with lower nutrient quality, recovery times remain similar among cells that do recover. Overall, since recovery from drug exposure is essentially a process of threshold crossing, stochasticity in the expression of resistance results in faster recovery than what is predicted by the deterministic model under most growth conditions.

## Discussion

Our model of the classical tetracycline response in *E. coli* combines biochemical reaction dynamics with gene regulation and global effects of the drug on metabolism to provide a framework to understand microbial antibiotic resistance in the context of dynamic and heterogeneous populations. Upon drug exposure, expression of resistance does not only depend on direct regulation by transcription factors but also on global effects on protein expression linked to the metabolic state of the cell [17,41,51]. And since restoration of metabolic functions (i.e., cell growth) also depends on the expression of resistance, this metabolism-mediated link introduces an additional feedback mechanism in the control of drug responses: failure to quickly deploy resistance genes results in higher levels of intracellular drug, leading to further reduction in expression of resistance and further drug accumulation. This positive feedback mediated by metabolic effects can act as a switch and is known to generate bistability and the coexistence of growing and arrested cells in the presence of antibiotics [18,52]. We note that this model does not include cell death explicitly, which is consistent with tetracycline being a bacteriostatic drug. Rather, when the intracellular drug concentration is too high, the cell is permanently arrested in a state with no cell growth or gene expression, which can be reversed if the drug challenge is relieved. Therefore, by considering the interplay between drug action, cell growth and expression of resistance, we are able to build a comprehensive model that explains the progression of antibiotic responses in single cells and, at the same time, captures the emergence of heterogeneity in the populations and colony-level collective behaviors.

Our models were able to recapitulate the complex dynamics observed experimentally in two different microfluidic experiments. A single-cell microfluidic experiment measuring phenotypic diversity during antibiotic responses found the emergence of remarkable diversity of growth and gene expression within isogenic populations of *tet* resistant *E. coli* strains [6]. These experiments observed the coexistence of three phenotypes: fast-growing recovered cells, arrested cells with little TetA expression, and temporary moribund cells that grow slowly and eventually either recover or arrest. Our deterministic model was able to generate all three phenotypes observed in single-cell responses, including the existence of the semi-stable low-growth moribund state, from where the system can escape to recovery if TetA concentrations are high enough. Simulations of our model suggest the existence of distinct stable phenotypic states, suggesting future analysis to determine the exact nature of these phenotypes, and particularly what differentiates the intriguing moribund state from complete arrest.

Our stochastic model recapitulates the emergence of phenotypic heterogeneity during drug responses, which is observed even in isogenic populations in homogeneous environments, explaining the coexistence of stable states corresponding to recovered and arrested cells. Gene expression in bacteria is known to be noisy, with large cell-to-cell variation, particularly in regulated genes. Stochastic models capture this variability, describing how naturally occurring noise in cellular processes is propagated through mechanisms of regulation to generate different outcomes during antibiotic responses. As observed experimentally, our simulations find that faltering expression of resistance upon exposure results in slow growth and delayed recovery, with large variation of recovery times. We find that at large drug concentrations, while the majority of the microbial population is arrested upon exposure, small subpopulations can still survive and later regenerate the population. Moreover, the model can quantitatively predict how environmental factors such as the drug dose or the nutritional quality of the medium determine the distribution of cell states. Therefore, our models can be used to determine the patterns of population-level growth and expression of resistance, which result from the sum of all single-cell phenotypes.

The framework developed with this specific model can be used as the basis to develop large-scale stochastic simulations of whole drug-resistance pathways, which will be useful in identifying new drug targets. However, while our full stochastic model is comprehensive, its simulations are slow due to the necessity of serially executing reactions spanning a wide range of timescales. Therefore, the model is limited in the number of molecules that can be simulated in tractable time, especially with regards to the ribosome pool. Future implementations of this model could use more sophisticated versions of the Gillespie algorithm such as tau leaping to reintroduce stochasticity from fast reactions that is lost with the adiabatic approximations [53,54]. Increasing the efficiency of the simulations allows the consideration of larger systems with more realistic numbers of components, resulting in more accurate quantitative predictions.

A second microfluidic experiment [7] characterizing how expression of resistance is coordinated during antibiotic responses across structured microbial populations (biofilms), where consumption of nutrients by cells closer to the surface generates chemical gradients towards the interior of the colony [7,55–57]. This experiment found that the quick arrest of fast-growing cells near the surface of the biofilm causes a redistribution of nutrients towards the interior and reactivates previously dormant cells at deeper layers (Figure S5). Because lower metabolism increases resistance levels, reactivated dormant cells can survive exposure and repopulate the biofilm, maintaining population-level growth even at drug concentrations high enough to kill whole planktonic populations. Interestingly, the same three phenotypes from the single-cell studies were observed: fast-growing cells with moderate TetA expression close to the surface of the biofilm, slow-growing cells with high TetA expression further into the biofilm, and arrested cells with little TetA expression in the interior. By considering the effects of nutrient conditions in cell growth and gene expression, our model was able to recapitulate the dynamics of spatially heterogeneous growth patterns and expression of resistance – most notably the reactivation of dormant cells upon drug exposure and the persistent accumulation of resistance in the dormant subpopulation. This serves as evidence that the ingredients of our model and their mutual coupling are indeed sufficient to explain the complex dynamics observed experimentally.

We find that slow-growing cells express higher levels of resistance genes and are better suited to survive sudden exposures. The dynamics of drug responses is characterized by markedly different gene expression levels between fast- and slow-growing cells. Even in the absence of drug, TetA and TetR expression were experimentally determined to decrease linearly with the cell growth rate, which is predicted by the theory of proteome partition (Figure 2cd). In poorer media, slow-growing cells are able to grow in the presence of the drug because of their higher expression of resistance. Therefore, the nutrient condition of the medium surrounding the cell determines its resistance profile. This is particularly relevant for biofilms, where nutrient gradients from the surface towards the interior of the colony generate an array of metabolic states that underlie collective resistance [58–66].

Microbial drug resistance in the real world is often at odds with lab measurements, with infections often returning from remission. Our model bridges the gap between the dynamics of drug responses at the single-cell level and the resulting collective behavior of the population, and helps to understand how subpopulations of microbial cells are able to resist exposure to high drug doses and regenerate the colony. A quantitative description of how cell responses are regulated in complex environments is crucial to understand community-level behaviors such as antibiotic resistance, pathogenesis, and biofilms, which often can be explained without invoking additional specialized mechanisms.

## Data Availability Statement

The original contributions presented in the study are included in the article/Supplementary Information and at our GitHub page: https://github.com/schultz-lab/Phys-Biol-2023. Further inquiries can be directed to the corresponding authors.

## Author Contributions

MS, DS, PB, and JPTC designed the mathematical model and wrote the manuscript. JPTC and MS implemented the model and ran the simulations. All authors contributed to the article and approved the submitted version.

## References

[1] Grkovic S, Brown M H and Skurray R A 2001 Transcriptional regulation of multidrug efflux pumps in bacteria Semin. Cell Dev. Biol. 12 225–37

[2] Florence D, Isabelle P, Roland L, Ekkehard C and Patrice C 2007 Modes and Modulations of Antibiotic Resistance Gene Expression Clin. Microbiol. Rev. 20 79–114

[3] Maheshri N and O’Shea E K 2007 Living with Noisy Genes: How Cells Function Reliably with Inherent Variability in Gene Expression Annu. Rev. Biophys. Biomol. Struct. 36 413–34

[4] Kærn M, Elston T C, Blake W J and Collins J J 2005 Stochasticity in gene expression: from theories to phenotypes Nat. Rev. Genet. 6 451–64

[5] Raj A and van Oudenaarden A 2008 Nature, Nurture, or Chance: Stochastic Gene Expression and Its Consequences Cell 135 216–26

[6] Schultz D, Palmer A C and Kishony R 2017 Regulatory Dynamics Determine Cell Fate following Abrupt Antibiotic Exposure Cell Syst. 5 509–517.e3

[7] Stevanovic M, Boukéké-Lesplulier T, Hupe L, Hasty J, Bittihn, P, and Schultz D 2022 Nutrient Gradients Mediate Complex Colony-Level Antibiotic Responses in Structured Microbial Populations Front. Microbiol. 13 740259

[8] Dewachter L, Fauvart M and Michiels J 2019 Bacterial Heterogeneity and Antibiotic Survival: Understanding and Combatting Persistence and Heteroresistance Mol. Cell 76 255–67

[9] Ackermann M 2015 A functional perspective on phenotypic heterogeneity in microorganisms Nat. Rev. Microbiol. 13 497–508

[10] Dhar N and McKinney J D 2007 Microbial phenotypic heterogeneity and antibiotic tolerance Curr. Opin. Microbiol. 10 30–8

[11] Vega N M and Gore J 2014 Collective antibiotic resistance: mechanisms and implications Curr. Opin. Microbiol. 21 28–34

[12] Martins B M C and Locke J C W 2015 Microbial individuality: how single-cell heterogeneity enables population level strategies Curr. Opin. Microbiol. 24 104–12

[13] Bhusal R P, Barr J J and Subedi D 2022 A metabolic perspective into antimicrobial tolerance and resistance The Lancet Microbe 3 e160–1

[14] Stokes J M, Lopatkin A J, Lobritz M A and Collins J J 2019 Bacterial Metabolism and Antibiotic Efficacy Cell Metab. 30 251–9

[15] Martínez J L and Rojo F 2011 Metabolic regulation of antibiotic resistance FEMS Microbiol. Rev. 35 768–89

[16] Klumpp S and Hwa T 2008 Growth-rate-dependent partitioning of RNA polymerases in bacteria Proc. Natl. Acad. Sci. 105 20245–50

[17] Klumpp S, Zhang Z and Hwa T 2009 Growth Rate-Dependent Global Effects on Gene Expression in Bacteria Cell 139 1366–75

[18] Deris J B, Kim M, Zhang Z, Okano H, Hermsen R, Groisman A and Hwa T 2013 The innate growth bistability and fitness landscapes of antibiotic-resistant bacteria Science 342 1237435

[19] Elowitz M B, Levine A J, Siggia E D and Swain P S 2002 Stochastic gene expression in a single cell Science 297 1183–6

[20] Eldar A and Elowitz M B 2010 Functional roles for noise in genetic circuits Nature 467 167–73

[21] Schultz D, Wolynes P G, Jacob E B and Onuchic J N 2009 Deciding fate in adverse times: Sporulation and competence in Bacillus subtilis Proc. Natl. Acad. Sci. 106 21027–21034

[22] Schultz D, Walczak A M, Onuchic J N and Wolynes P G 2008 Extinction and resurrection in gene networks Proc. Natl. Acad. Sci. 105 19165–70

[23] Schultz D, Onuchic J N and Wolynes P G 2007 Understanding stochastic simulations of the smallest genetic networks J. Chem. Phys. 126 245102

[24] Meier I, Wray L V and Hillen W 1988 Differential regulation of the Tn10-encoded tetracycline resistance genes tetA and tetR by the tandem tet operators O1 and O2. EMBO J. 7 567–72

[25] Schultz D, Jacob E B, Onuchic J N and Wolynes P G 2007 Molecular level stochastic model for competence cycles in Bacillus subtilis Proc. Natl. Acad. Sci. 104 17582–7

[26] Kussell E and Leibler S 2005 Phenotypic Diversity, Population Growth, and Information in Fluctuating Environments Science 309 2075–8

[27] Süel G M, Garcia-Ojalvo J, Liberman L M and Elowitz M B 2006 An excitable gene regulatory circuit induces transient cellular differentiation Nature 440 545–50

[28] Ozbudak E M, Thattai M, Kurtser I, Grossman A D and Van Oudenaarden A 2002 Regulation of noise in the expression of a single gene Nat. Genet. 31 69–73

[29] Fernández L and Hancock R E W 2012 Adaptive and mutational resistance: Role of porins and efflux pumps in drug resistance Clin. Microbiol. Rev. 25 661–81

[30] Le T T, Harlepp S, Guet C C, Dittmar K, Emonet T, Pan T and Cluzel P 2005 Real-time RNA profiling within a single bacterium Proc. Natl. Acad. Sci. 102 9160–9164

[31] Le T T, Emonet T, Harlepp S, Guet C C, Cluzel P 2006 Dynamical determinants of drug-inducible gene expression in a single bacterium Biophys J 90 3315–21

[32] Muthukrishnan A-B, Kandhavelu M, Lloyd-Price J, Kudasov F, Chowdhury S, Yli-Harja O and Ribeiro A S 2012 Dynamics of transcription driven by the tetA promoter, one event at a time, in live Escherichia coli cells Nucleic Acids Res. 40 8472–83

[33] Ramos J L, Martínez-Bueno M, Molina-Henares A J, Terán W, Watanabe K, Zhang X, Gallegos M T, Brennan R and Tobes R 2005 The TetR Family of Transcriptional Repressors Microbiol. Mol. Biol. Rev. 69 326–56

[34] Eckert B and Beck C F 1989 Overproduction of transposon Tn10-encoded tetracycline resistance protein results in cell death and loss of membrane potential J. Bacteriol. 171 3557–9

[35] Carvalho G, Forestier C and Mathias J D 2019 Antibiotic resilience: a necessary concept to complement antibiotic resistance? Proc. R. Soc. B Biol. Sci. 286 20192408

[36] Moran M A, Satinsky B, Gifford S M, Luo H, Rivers A, Chan L K, Meng J, Durham B P, Shen C, Varaljay V A, Smith C B, Yager P L and Hopkinson B M 2013 Sizing up metatranscriptomics ISME J. 7 237–43

[37] Gillespie D T 1976 A general method for numerically simulating the stochastic time evolution of coupled chemical reactions J. Comput. Phys. 22 403–34

[38] Schnappinger D and Hillen W 1996 Tetracyclines: antibiotic action, uptake, and resistance mechanisms Arch. Microbiol. 165 359–69

[39] Chopra I and Roberts M 2001 Tetracycline antibiotics: mode of action, applications, molecular biology, and epidemiology of bacterial resistance Microbiol. Mol. Biol. Rev. 65 232–60

[40] Scott M, Gunderson C W, Mateescu E M, Zhang Z and Hwa T 2010 Interdependence of cell growth and gene expression: Origins and consequences Science 330 1099–102

[41] Klumpp S and Hwa T 2014 Bacterial growth: global effects on gene expression, growth feedback and proteome partition Curr. Opin. Biotechnol. 28 96–102

[42] Camas F M, Bláquez J and Poyatos J F 2006 Autogenous and nonautogenous control of response in a genetic network Proc. Natl. Acad. Sci. 103 12718–23

[43] Rosenfeld N, Elowitz M B and Alon U 2002 Negative Autoregulation Speeds the Response Times of Transcription Networks J Mol Biol 323 785–93

[44] Dublanche Y, Michalodimitrakis K, Kümmerer N, Foglierini M and Serrano L 2006 Noise in transcription negative feedback loops: simulation and experimental analysis Mol. Syst. Biol. 1–12

[45] Madar D, Dekel E, Bren A and Alon U 2011 Negative auto-regulation increases the input dynamic-range of the arabinose system of Escherichia coli BMC Syst. Biol. 5 111

[46] Nevozhay D, Adams R M and Murphy K F 2009 Negative autoregulation linearizes the dose – response and suppresses the heterogeneity of gene expression Proc. Natl. Acad. Sci. 106 5123–8

[47] Sigler A, Schubert P, Hillen W and Niederweis M 2000 Permeation of tetracyclines through membranes of liposomes and Escherichia coli Eur. J. Biochem. 267 527–34

[48] Reuter A, Virolle C, Goldlust K, Berne-Dedieu A, Nolivos S and Lesterlin C 2020 Direct visualisation of drug-efflux in live Escherichia coli cells FEMS Microbiol. Rev. 44 782–92

[49] Wang P, Robert L, Pelletier J, Dang W L, Taddei F, Wright A and Jun S 2010 Robust Growth of Escherichia coli Curr. Biol. 20 1099–103

[50] Wald-Dickler N, Holtom P and Spellberg B 2018 Busting the Myth of “Static vs Cidal”: A Systemic Literature Review Clin. Infect. Dis. 66 1470–4

[51] Greulich P, Scott M, Evans M R and Allen R J 2015 Growth-dependent bacterial susceptibility to ribosome-targeting antibiotics Mol. Syst. Biol. 11 796

[52] Frenkel N, Dover R S, Titon E, Shai Y and Rom-Kedar V 2021 Bistable Bacterial Growth Dynamics in the Presence of Antimicrobial Agents Antibiotics 10

[53] Cao Y, Gillespie D T and Petzold L R 2005 Avoiding negative populations in explicit Poisson tau-leaping J. Chem. Phys. 123 54104

[54] Gillespie D T 2007 Stochastic Simulation of Chemical Kinetics Annu. Rev. Phys. Chem. 58 35–55

[55] Bittihn P, Didovyk A, Tsimring L S and Hasty J 2020 Genetically engineered control of phenotypic structure in microbial colonies Nat. Microbiol. 5 697–705

[56] Flemming H C, Wingender J, Szewzyk U, Steinberg P, Rice S A and Kjelleberg S 2016 Biofilms: an emergent form of bacterial life Nat. Rev. Microbiol. 14 563–75

[57] Jeckelmann J M and Erni B 2020 Transporters of glucose and other carbohydrates in bacteria Pflügers Arch. - Eur. J. Physiol. 472 1129–53

[58] Stewart P S and Franklin M J 2008 Physiological heterogeneity in biofilms Nat. Rev. Microbiol. 6 199–210

[59] Besharova O, Suchanek V M, Hartmann R, Drescher K and Sourjik V 2016 Diversification of gene expression during formation of static submerged biofilms by Escherichia coli Front. Microbiol. 7 1568

[60] Cao Y, Ryser M D, Payne S, Li B, Rao C V and You L 2016 Collective Space-Sensing Coordinates Pattern Scaling in Engineered Bacteria Cell 165 620–30

[61] Bottery M J, Pitchford J W and Friman V-P 2021 Ecology and evolution of antimicrobial resistance in bacterial communities ISME J. 15 939–48

[62] Orazi G and O’Toole G A 2019 “It Takes a Village”: Mechanisms Underlying Antimicrobial Recalcitrance of Polymicrobial Biofilms J. Bacteriol. 202 e00530–19

[63] Kowalski C H, Morelli K A, Schultz D, Nadell C D and Cramer R A 2020 Fungal biofilm architecture produces hypoxic microenvironments that drive antifungal resistance Proc. Natl. Acad. Sci. 117 22473–22483

[64] MacLean R C, Hall A R, Perron G G and Buckling A 2010 The population genetics of antibiotic resistance: integrating molecular mechanisms and treatment contexts Nat. Rev. Genet. 11 405–14

[65] Bennett B D, Kimball E H, Gao M, Osterhout R, Van Dien S J and Rabinowitz J D 2009 Absolute metabolite concentrations and implied enzyme active site occupancy in Escherichia coli Nat. Chem. Biol. 5 593–9

[66] Kim J, Darlington A, Salvador M, Utrilla J and Jiménez J I 2020 Trade-offs between gene expression, growth and phenotypic diversity in microbial populations Curr. Opin. Biotechnol. 62 29–37

