## Supplementary Information for "Dynamical model of antibiotic responses linking expression of resistance to metabolism explains emergence of heterogeneity during drug exposures"

Table S1. Definitions, values, and interpretations of parameters and variables used in deterministic simulations.

| Symbol | Value/Expression | Interpretation | Symbol | Value/Expr. | Interpretation |
|---|---|---|---|---|---|
| $K_i$ | $0.015\ min^{-1}$ | drug diffusion rate | $K_b$ | $60\ min^{-1}$ | dissociation constant for TetR and operator |
| $D(t)$ | varies | extracellular drug concentration | $A$ | $0.008\ \mu M min^{-1}$ | maximal TetA promoter activity |
| $K_a$ | $50\ min^{-1}$ | catalytic rate constant of TetA | $r_{0a}$ | $0.0001\ \mu M$ | dissociation constant for free TetR and its DNA binding site |
| $d_f$ | $\frac{1}{2}\left(d - r - K_d + \sqrt{(d - r - K_d)^2 + 4K_d d}\right)$ | free intracellular drug | $H_r(r_f)$ | $R\frac{r_{0,r}^4}{r_{0,r}^4 + r_f^4}$ | TetR synthesis |
| $k_a$ | $10\ \mu M$ | Michaelis constant for TetA | $R$ | $0.0003\ \mu M min^{-1}$ | maximal TetR promoter activity |
| $\lambda_0$ | $0.015\ min^{-1}$ or $0.029\ min^{-1}$ | growth at no drug, full nutrients | $r_{0r}$ | $0.000075\ \mu M$ | dissociation constant for free TetR and its DNA binding site |
| $\kappa_t$ | $\kappa_t^0 \frac{K_{ribo}}{K_{ribo} + d_f}$ | translational capacity | $\phi_c$ | 0.48 | constant |
| $\kappa_t^0$ | $4.5\ h^{-1}$ | translational capacity at no drug, full nutrients | $\rho$ | 0.76 | constant |
| $\kappa_n$ | $0.035\ min^{-1}$ or $0.1196\ min^{-1}$ | nutritional capacity | $K_d$ | $0.001\ \mu M$ | dissociation constant for TetR and drug |
| $H_a(r_f)$ | $A\frac{r_{0,a}^4}{r_{0,a}^4 + r_f^4}$ | TetA synthesis | $RR$ | $N_r \lambda_0 \frac{\kappa_n}{\kappa_n + \kappa_t^0}$ | number of free ribosomes in the cell |
| $r_f$ | $\frac{1}{2}\left(r - d - K_d + \sqrt{(r - d - K_d)^2 + 4K_d r}\right)$ | free TetR | $N_R^{max}$ | 100 | number of ribosomes in the cell |
| $g$ | $f\frac{\kappa_t^0}{\kappa_t}$ | dependence of ribosome synthesis on translation inhibition and nutrient levels | $K_{ribo}$ | $1\ \mu M$ | dissociation constant for drug and ribosome |

**Nondimensional Model**
We rescale time and reaction rates by the maximum growth rate $\lambda_0$, intracellular drug by its dissociation constant for ribosome unbinding, repressor concentration by its dissociation constant for drug unbinding, and pump concentration by its catalytic rate. All rescaled variables and parameters in our model are summarized in Table S2. The resulting nondimensional system, shown below (primes dropped for simplicity), has fewer parameters, is normalized, and allows us to better understand the order of magnitude of values in the model:

$$\dot{d} = K_i\left(D(t) - d_f\right) - \frac{a d_f}{1 + K_1 d_f} - \lambda d$$
$$\dot{a} = f\ H_a\left(r_f\right) - \lambda a$$
$$\dot{r} = f\ H_r\left(r_f\right) - \lambda r$$

Table S2. Definitions, values, and interpretations of parameters and variables used in the nondimensional model.

| Symbol | Expr. | Interpretation | Symbol | Expression | Interpretation |
|---|---|---|---|---|---|
| $t'$ | $\lambda_0 t$ | time scaled by growth rate | $R'$ | $\frac{R}{\lambda_0 K_d}$ | maximal TetR promoter activity scaled by maximal growth rate and the drug-TetR dissociation constant |
| $d'$ | $\frac{d}{K_{ribo}}$ | intracellular drug concentration relative to the ribosome dissociation constant | $A'$ | $\frac{AK_a}{k_a \lambda_0^2}$ | maximal TetA promoter activity scaled by maximal growth rate and the Michaelis constant for TetA |
| $r'$ | $\frac{r}{K_d}$ | repressor concentration relative to the drug-TetR dissociation constant | $r'_{0a}$ | $\frac{r_{oa}}{K_d}$ | dissociation constant for free TetR and its TetA DNA binding site relative to the drug-TetR dissociation constant |
| $a'$ | $\frac{K_a}{k_a \lambda_0} a$ | pump concentration relative to its catalytic rate | $r'_{0r}$ | $\frac{r_{or}}{K_d}$ | dissociation constant for free TetR and its TetR DNA binding site relative to the drug-TetR dissociation constant |

| | | | | | |
|---|---|---|---|---|---|
| $\kappa_t'$ | $\frac{1}{1+{d_f}'}$ | translational capacity | $H_r'(H_f')$ | $\frac{R'}{1+\left(\frac{r_f}{{r_{o,r}}'}\right)^4}$ | TetR synthesis |
| $\lambda'$ | $\frac{\lambda}{\lambda_0}$ | growth rate relative to maximum growth rate | $f'$ | $\frac{\lambda'^2}{\kappa_n}$ | dependence of protein synthesis on translation inhibition and nutrient levels |
| $K_i'$ | $K_i/\lambda_0$ | drug diffusion rate scaled by maximum growth rate | $d_f'$ | $\frac{1}{2K_2}\Big(K_2 d - r - 1 + \sqrt{(K_2 d - r - 1)^2 + 4K_2 d}\Big)$ | free intracellular drug |
| $K_1$ | $K_{ribo}/k_a$ | dissociation constant for drug and ribosome relative to the Michaelis constant for TetA | $r_f'$ | $\frac{1}{2}\Big(r - K_2 d - 1 + \sqrt{(r - K_2 d - 1)^2 + 4r}\Big)$ | free TetR |
| $K_2$ | $K_{ribo}/K_d$ | dissociation constant for drug and ribosome relative to the dissociation constant for TetR and drug | $K_x$ | $\frac{\kappa_n}{\kappa_t^0}$ | maximum nutritional capacity relative to the maximum translational capacity |
| $H_a'(H_f')$ | $\frac{A'}{1+\left(\frac{r_f}{{r_{o,a}}'}\right)^4}$ | TetA synthesis | | | |

**The effects of the drug on the proteome partition are necessary to explain the advantage of slow-growing cells**

Here we simulate the drug response assuming that TetA and TetR belong to the Q sector of proteins that are not affected by the metabolic state of the cell or by translation inhibition. In this case, the expression for the modulation of protein expression $f = (\lambda/\lambda_0)\ [\kappa_t/(\kappa_t + \kappa_n)]\ /\ [\kappa_t^0/(\kappa_t^0 + \kappa_n)]$ reduces to $f = \lambda/\lambda_0$, as assuming that the nutritional and translational capacities of the cell do not affect the expression of TetA and TetR means we can set $\kappa_t = \kappa_t^0$.

When using this expression for $f$ in our simulations, the advantage of slow-growing cells disappears (Figure S1). Under the assumption that TetR and TetA are Q-sector proteins, fast-growing cells have the advantage of diluting drug faster in the cytoplasm, while drug influx does not change. They therefore have higher growth rates than slow-growing cells at the end of the response for the same drug concentration.

**Nutritional capacity in our simulations was determined from the maximum growth rate in our experiments**

In the single-cell microfluidic experiment, minimal M63 medium was used, whereas in the microcolony microfluidic experiment, EZ Rich defined medium was used. The maximum growth rate was 0.015/min in the former, and 0.029/min in the latter. The $\kappa_n$ in our simulations was calculated using the formula $\lambda_0 = (\phi_c/\rho) \cdot \kappa_t^0\ \kappa_n/(\kappa_t^0 + \kappa_n)$, where $\kappa_t^0$ is 4.5 $h^{-1}$ and $\lambda_0$ is the maximum growth rate in our experiments. In Figure 1, $\lambda_0$ was 0.015/min, and in Figure 2, $\lambda_0$ was 0.029/min.

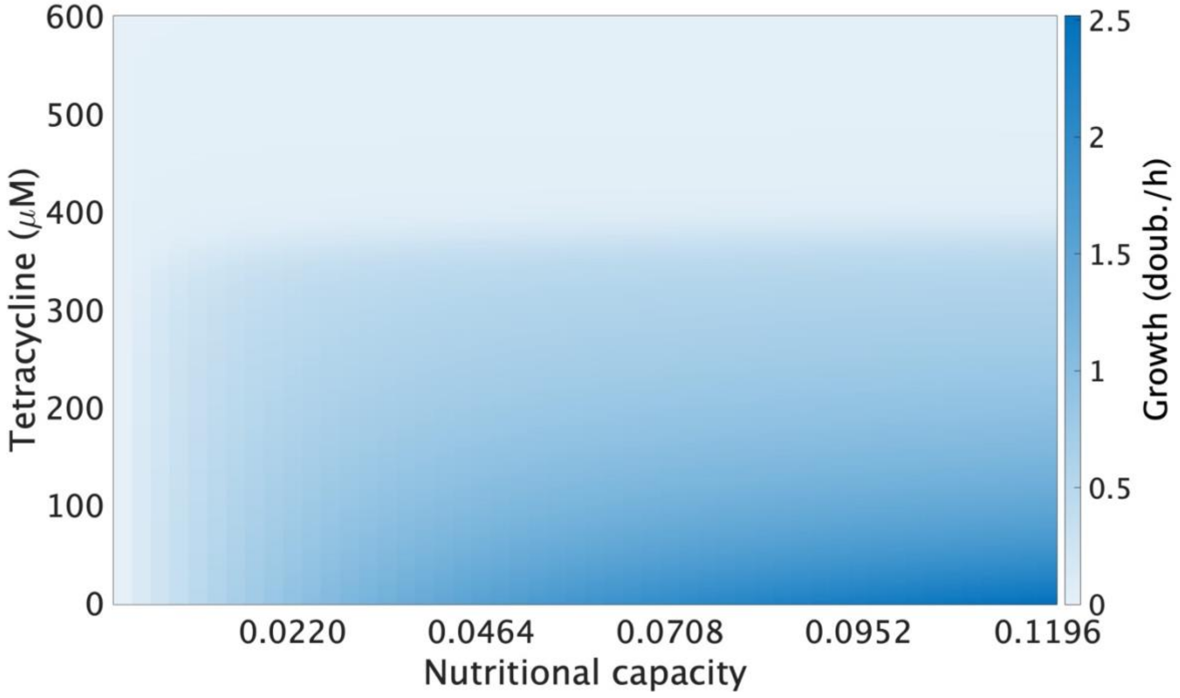


**Figure S1. Slow-growing cells do not have an advantage over fast growing ones if TetA and TetR are not affected by nutritional and translational inhibition.** Growth rates at different combinations of extracellular drug concentrations and nutritional capacities where TetA and TetR are modeled as belonging to the Q sector.

**Modeling TetR as a Dimer in the Binding of the Operators in the Stochastic Model**
As described in the main text, we model the gene expression of TetR and TetA using Hill functions. The Hill coefficient is one of its parameters, and in general, it reflects the cooperativity of ligand molecules, which in the case of our model are TetR molecules. As such, the Hill coefficient also determines the steepness of the Hill function as the number of ligands required to bind to the receptor in order to elicit a functional effect, in this case gene expression, determines how close to the step-function the Hill function is. In the dynamical model, we set the Hill coefficient of TetR to 4 when modeling the expression of TetR, and we also set it to 4 when modeling the expression of TetA. We determined these Hill coefficients by fitting Hill functions to our experimental data [6]. In the stochastic model, on the other hand, we model the action of TetR as that of a dimer. We do so in order to both match the deterministic model and to account for the regulatory architecture of the *tet* operon (Figure S2). The regulatory region of the operon consists of two operators to which TetR binds, and three promoters to which RNA polymerase can bind. Two of the promoters, $P_{R1}$ and $P_{R2}$ regulate transcription initiation for TetR and the remaining one, $P_A$, regulates transcription initiation for TetA.

Note that in order for RNA polymerase to bind to $P_A$ and transcribe the gene for TetA, both operators $O_1$ and $O_2$ have to be free. Therefore, the rate of TetA production is proportional to the probability that $O_1$ is unbound *times* the probability that $O_2$ is unbound. If we model the binding of TetR and $O_1$ by a Hill function, and the binding of TetR and $O_2$ by another Hill function, the production of TetA is proportional to the product of these two functions. Therefore, modeling TetR as a dimer is equivalent to both of these Hill functions having a Hill coefficient of 2, and so their product is a Hill function with coefficient 4, which matches closely the empirically determined Hill coefficient of 4 for the expression of TetA (Figure S3).

On the other hand, even if $O_2$ is bound to TetR, as long as $O_1$ if free, RNA polymerase can bind to $P_{R1}$ and transcribe the gene for TetR. As a result, the rate of TetR production is proportional to the probability that $O_1$ is unbound *plus* the probability that both $O_1$ and $O_2$ are unbound. If we model TetR as a dimer, the latter, as we already showed in the paragraph above, is a Hill function with coefficient 4, and the former is simply a Hill function with coefficient 2. The sum of these two Hill functions can reasonably be approximated by a Hill function with a Hill coefficient of 4 that we have in the deterministic model (Figure S3).

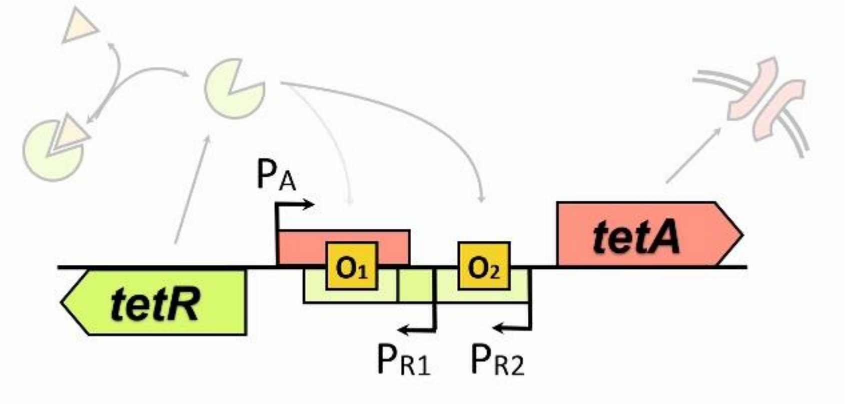


**Figure S2. The Configuration of the *tet* Regulatory Region.** The operators are shown as yellow squares, and the promoters are shown as arrows. $P_A$ represents the promoter for TetA, and $P_{R1}$ and $P_{R2}$ represent the promoters for TetR.

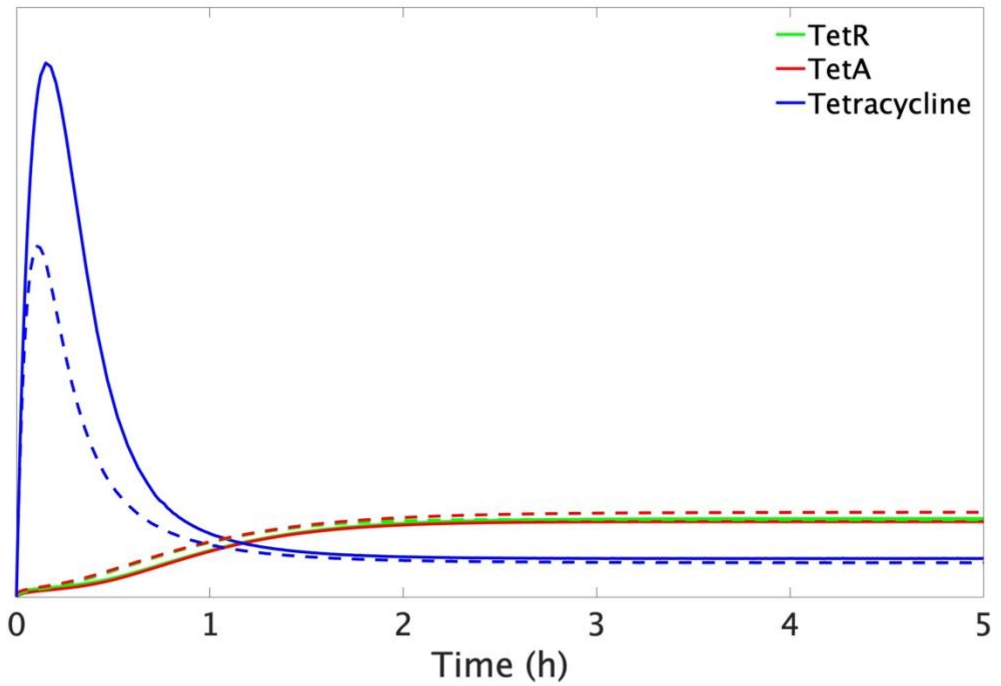


**Figure S3. Response dynamics is not sensitive to Hill coefficient**. Cell responses to sudden drug exposure calculated with the deterministic model using (i) single Hill functions for the regulation of TetA and TetR, both with coefficient 4 (dashed lines), and (ii) combinations of Hill functions, one for each TetR binding site interfering with expression, reflecting the true architecture of the *tet* promoter (solid lines). Each Hill function has a coefficient of 2, reflecting the binding of TetR dimers, so the multiplication of two Hill functions also effectively has a coefficient of 4.

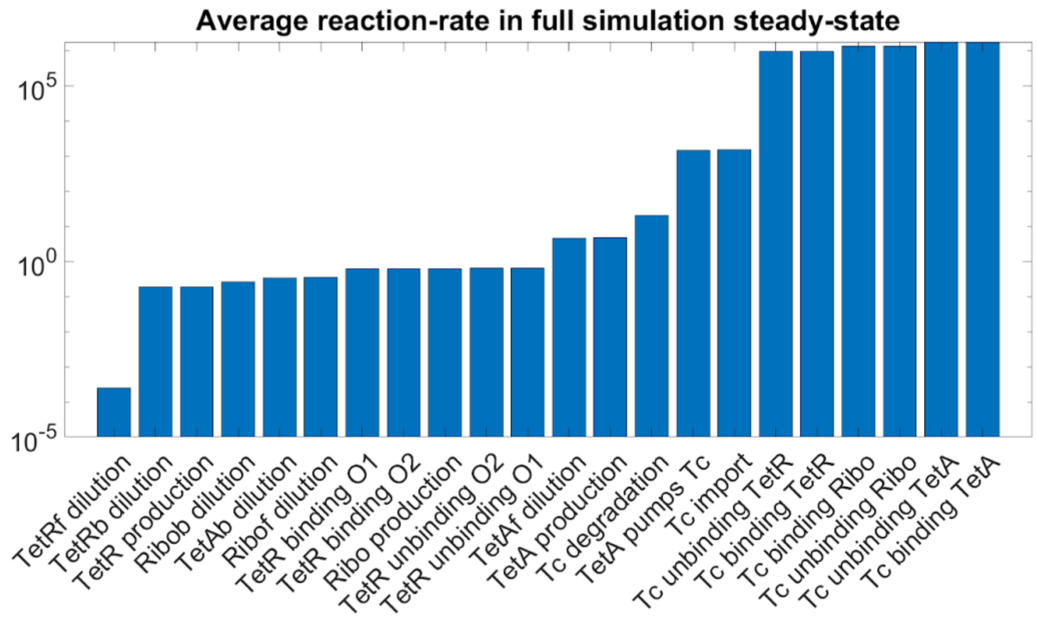


**Figure S4. Distribution of reaction times in the stochastic model where all reactions are simulated stochastically.** In the simulation where all 22 reactions are computed, high tetracycline (Tc) concentrations make reactions where Tc binds any molecule have much higher reaction rates than others. This manifests as reactions that are much more likely to occur, quickly reaching chemical equilibrium.

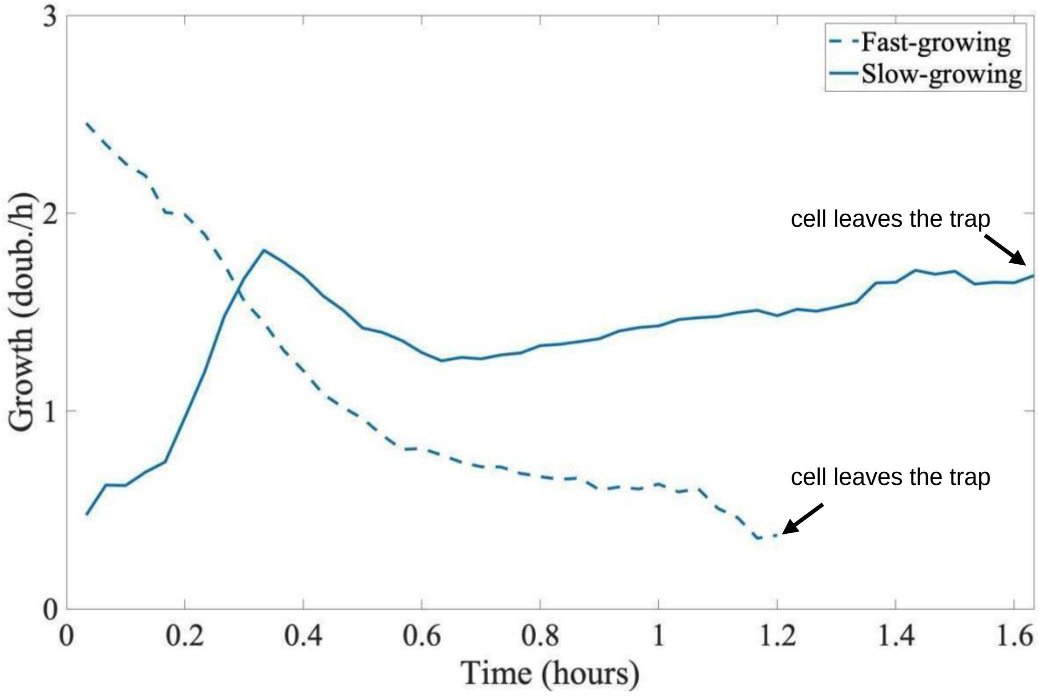


**Figure S5. Growth upon drug exposure along the trajectories of two cells in the microfluidic trap.** A cell that is growing fast at the top of the trap prior to drug exposure experiences significant growth reduction once the drug is introduced (at time zero). A slow-growing cell from the interior of the colony survives exposure and is reactivated by the increased supply of nutrients diffusing down the trap, increasing growth before being pushed out of the trap.
